# CoCoCoNet: Conserved and Comparative Co-expression Across a Diverse Set of Species

**DOI:** 10.1101/2020.04.21.053900

**Authors:** John Lee, Manthan Shah, Sara Ballouz, Megan Crow, Jesse Gillis

## Abstract

Co-expression analysis has provided insight into gene function in organisms from Arabidopsis to Zebrafish. Comparison across species has the potential to enrich these results, for example by prioritizing among candidate human disease genes based on their network properties, or by finding alternative model systems where their co-expression is conserved. Here, we present CoCoCoNet as a tool for identifying <u>co</u>nserved gene modules and <u>co</u>mparing <u>co</u>-expression <u>net</u>works. CoCoCoNet is a resource for both data and methods, providing gold-standard networks and sophisticated tools for on-the-fly comparative analyses across 14 species. We show how CoCoCoNet can be used in two use cases. In the first, we demonstrate deep conservation of a nucleolus gene module across very divergent organisms, and in the second, we show how the heterogeneity of autism mechanisms in humans can be broken down by functional groups, and translated to model organisms. CoCoCoNet is free to use and available to all at https://milton.cshl.edu/CoCoCoNet, with data and R scripts available at ftp://milton.cshl.edu/data.

## INTRODUCTION

How a gene’s expression level changes across conditions is a rich source of information about its function, a fact which gene co-expression networks aim to capture in a general framework (1). Gene co-expression networks link genes by their similarity in expression pattern, yielding connected subnetworks which are likely to share biological functions (2). One of the most important uses of co-expression networks is to test whether a newly identified set of genes forms a clear module (3). Once that is established, the specific topology within the network can be studied in detail to determine central nodes or to define critical co-expression relationships (4, 5).

The utility of expression as a readout across biological systems has allowed co-expression network analysis to be applied very broadly: to group and classify genes in model organisms (e.g., Arabidopsis (6, 7), mice (8), and yeast (9)), to find and characterize disease genes (e.g., in autism (10), Parkinson’s disease (11), and heart disease (12)), and as an important contributor to sophisticated algorithms for inferring gene properties (e.g., miRNA targets (13), transcription factor regulation (14), and GO annotations (15, 16)). Because evolution often works by rewiring existing gene-gene relationships, a particularly important area of co-expression analysis is cross-species comparison. Though it is well-established that cross-species analyses can enrich for biologically relevant modules (17), even simple comparisons remain very challenging. With CoCoCoNet, we have aimed to systematize comparative co-expression, expanding the range of species covered in the field as a whole, improving the statistical rigor of network analysis within each species, and enhancing the sophistication of integrative analyses across species.

CoCoCoNet allows users to access novel research areas by querying and comparing well-powered co-expression networks for 14 species. With a few clicks, researchers can input their gene or genes of interest, and look for co-expression relationships that may be conserved across large phylogenetic distances. While co-expression is a key component of other web servers and databases such as COXPRESdb (18), ATTED-II (19), GeneFriends (20), PlaNet (21), MouseNet (22) and GeneMANIA (23), few provide data beyond the standard model organisms (human, mouse, fly, roundworm, yeast, and Arabidopsis). Those that do, lack the ability to make cross-species comparisons. For example, PlaNet, COXPRESdb and ATTED-II provide co-expression data for several of the species we cover, but there are no convenient methods to directly compare the networks, nor do they perform any explicit analyses of co-expression strength. In contrast, CoCoCoNet provides users with convenient access to both data and methods for cross-species analyses. This opens up a range of potential research questions, such as:

- Which genes are related to my target gene, and do those relationships change across species?
- When has co-expression been conserved across large phylogenetic distances?
- Does my gene set subdivide into clusters that are maintained across species?
- Is my gene of interest co-expressed with other genes of interest in species I do not study?

In the following, we summarize the methods, data, and operation of CoCoCoNet and walk through two use cases: one focused on highly co-expressed modules in yeast, and another on autism disease genes. In addition, we provide substantially expanded detail in our supplement – providing details on network construction, resources used, quality control, and a complete walk-through of the webserver. We have made all methods and data available for use by other researchers, including the underlying network data and methods for assessing it.

## MATERIAL AND METHODS

### RNA-seq datasets

Because the quality of co-expression data is highly correlated with the total number of samples across all datasets (4), we aimed to collect as much data as possible for each species. To this end, we searched NCBI’s Sequence Read Archive (SRA) database (24), using the R Bioconductor package “SRAdb” (25) for bulk RNA-sequencing datasets (unique SRA Study IDs), excluding those with fewer than 10 samples. Cancer-related studies were also excluded since they are not likely to generalize well. To maximize the independence of co-expression measurements within individual datasets, we included only one replicate (a.k.a. “run accession”) per unique Biosample ID, choosing the replicate with the maximum amount of data by number of spots. Reference genomes and genome annotation files were downloaded from ENSEMBL (26) (Sept 2019). Sequence reads were downloaded directly from NCBI’s ftp site (ftp://ftp.sra.ebi.ac.uk/vol1/fastq) and were aligned to the reference genome using STAR v2.6.0c (27). See table S1 for more details.

Datasets identified in SRAdb were included in our gold-standard co-expression networks if they met two additional criteria: measurable expression of at least 50% of all genes (figure S2); and above-threshold similarity to an aggregate expression profile characterizing all datasets. Procedurally, this means that for every sample, we rank genes by expression level, then average these ranks across all samples within a dataset, and finally average these dataset-level results to obtain a “global average”. Next, we compute the Spearman’s correlation between each sample in each dataset and this global average (figures S3 and S4). If the average of the worst 10 correlation coefficients is less than 0.3, we remove that dataset entirely.

**Figure 1.**
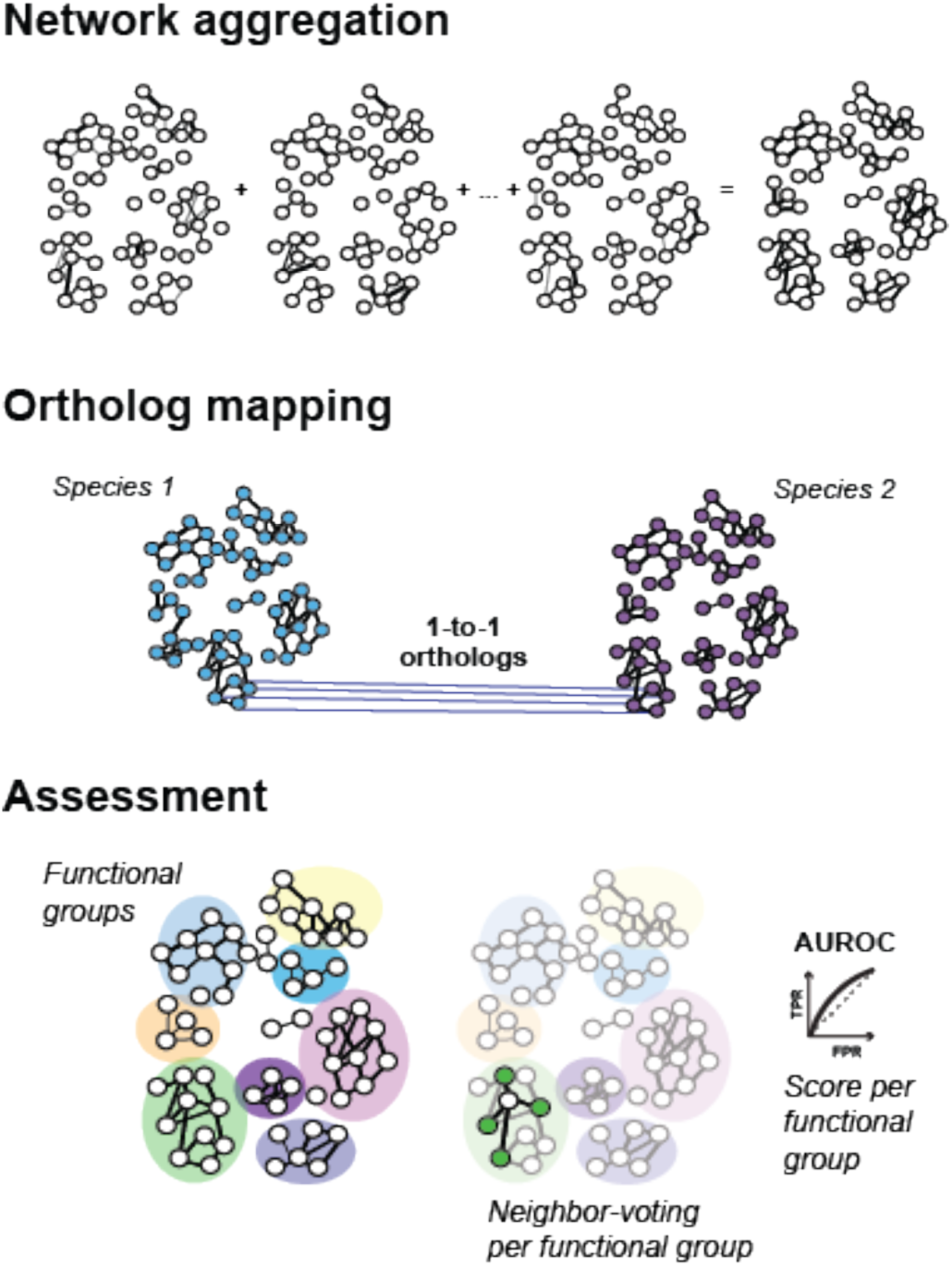
Schematic of underlying data. Co-expression networks are aggregated for each species, ortholog maps are generated for each pair of species, and data quality of data is assessed using a neighbor voting algorithm across all functional groups.

In combination with our minimal sample requirements, these checks ensure that each dataset used in the aggregation of our co-expression networks is both well-powered and likely to generalize. Figure 2 contains a detailed summary of the number of experiments and the total number of samples that went into the construction of the aggregate networks. Further detail on these datasets can be found in tables S2 and S4.

**Figure 2.**
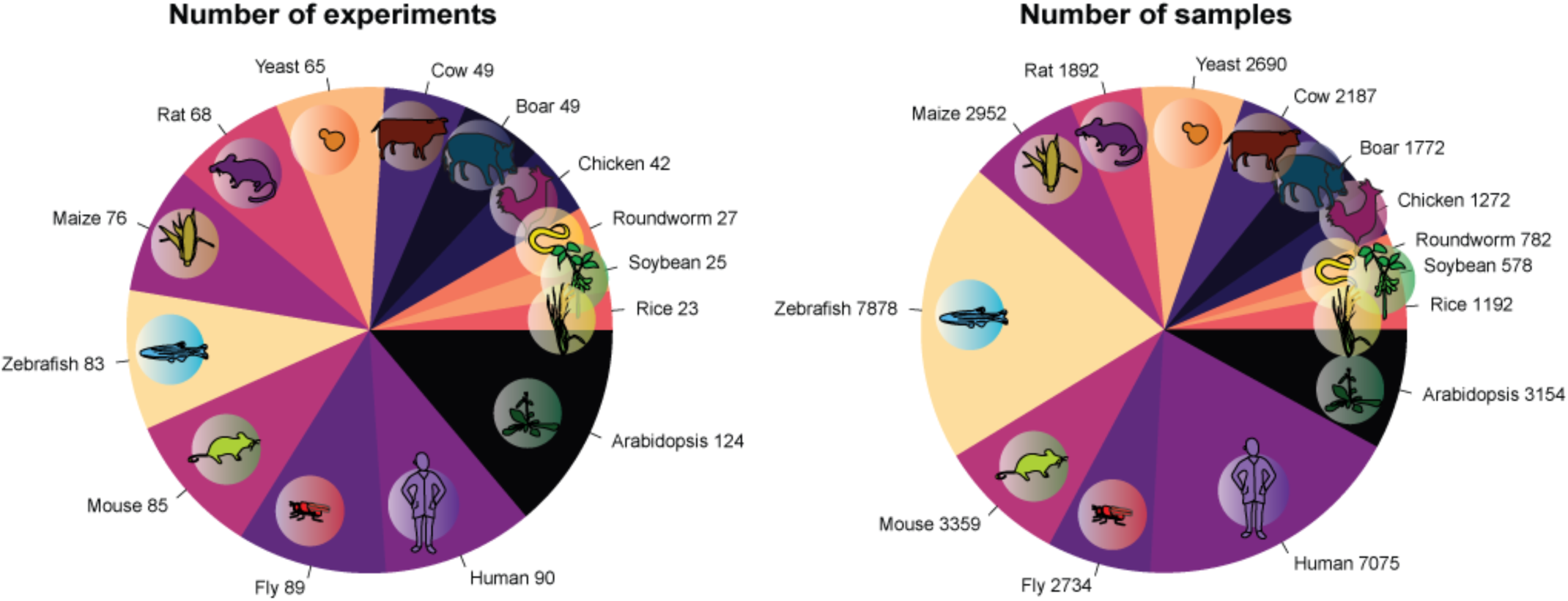
Left: Counts of experiments expressing at least half of all genes. Right: Counts of samples with a correlation with the global average greater than 0.3.

We note that we did not limit our search to a single sequencing platform. In general, platform consistency is maintained within experiments, and co-expression networks are independently constructed and standardized, thus the aggregation of these controlled networks is not affected by this class of variability. In total, our data comprises of 39,517 samples across the 14 species, 34,729 of which utilize Illumina HiSeq 2000 or 2500 (table S4).

### Co-expression network construction and aggregation

Co-expression networks for each dataset were constructed by computing Spearman’s correlation between every pair of genes (figure S1). This generates a network that is then rank standardized, and normalized by dividing through by the maximum rank (4). Genes that are not observed in a particular dataset naturally have no variance, making correlation computations impossible. We replace these NA values with the median value of the network. Networks obtained from individual datasets were then aggregated by adding all of the network adjacency matrices, then rank standardizing and dividing by the maximum rank (figure 1).

While other co-expression tools use Pearson’s correlation as their primary metric (18, 19, 20), we use Spearman’s correlation. We have shown in (4) that there is marginal difference in performance using Pearson’s correlation over Spearman’s Correlation. We utilize the non-parametric approach of Spearman’s to ensure that outlier values do not have undue influence, allowing results to be driven by the power of larger data.

Within CoCoCoNet, users can choose to query aggregates built with almost all genes, or those built with a smaller high-confidence set. Our minimal filter requires that genes be expressed at least once in at least half of the datasets. Genes that fail to meet this requirement are removed from the aggregate co-expression network, yielding the “almost all genes” set. A more stringent filter allows for faster processing, and provides greater confidence in retained links. To filter for genes that are well-powered, we count the number of datasets where a gene has at least 10 reads in each of 10 or more samples. “High confidence genes” are those that meet these criteria in more than 20 datasets.

### Gene annotations and ortholog mapping

We use the Gene Ontology (GO) (28, 29) to obtain gene function annotations. GO terms and gene associations were obtained by merging data from NCBI’s website (ftp://ftp.ncbi.nlm.nih.gov/gene/DATA/gene2go.gz) (Jan 2020) and the Bioconductor package “biomaRt” (30) (table S3). Terms were then propagated in the ontology tree using a transitive property and filtered to include terms annotating between 10 and 1000 genes. These are then used in enrichment analyses, performed using Fisher’s exact test followed by an FDR correction.

Ortholog data is obtained from OrthoDB (31), allowing us to provide 1-to-1 ortholog maps for every pair of species included in CoCoCoNet (table S5). This is accomplished by searching for the most recent phylogenetic split between the two query species, and obtaining inferred orthology groups for all genes descended from the common ancestor. Genes are then filtered to the corresponding input species and mapped to each other (figure 1).

### Network assessment

Guilt-by-association based methods are used to ascertain the quality of co-expression networks (32), and can also be used to determine the connectivity of a gene set. To accomplish this, CoCoCoNet implements functions from the Bioconductor package “EGAD” (33) on the gene set provided by the user, along with the orthologs from the second species selected, and GO annotations (figure 1). EGAD measures the performance of a network and a gene set through the neighbor-voting algorithm and reports an area under the receiver operating curve (AUROC) or the area under the precision recall curve (AUPRC). These performances can also be compared to predictions based solely on node degree (34). AUCs close to 0.5 indicate poor performance, 0.7 being quite good, and 1 being perfect. If the AUCs from both species are high, the tested gene sets and their co-expression modules can be said to be conserved, particularly if the node degree bias is low.

### Implementation

This web server is implemented using the open source R Shiny Server (35). In our networks, nodes are genes and edges are normalized average correlation statistics across all underlying datasets, as detailed above. Visual clustering of each network is implemented using the physical properties of the network and the “visNetwork” R package (36). We assign each node a mass proportional to its total node degree, where the larger the mass, the more repulsive the node. A Barnes-Hut n-body simulation (37) is applied, forcing high degree nodes towards cluster centers and low degree nodes towards the cluster peripheries. Network data is stored in HDF5 format, which allows for rapid search of specified data. Histograms and scatter plots are generated using the R package “ggplot2” (38) and made interactive using the R package “plotly” (39).

### Web server description

CoCoCoNet is designed to be simple to use and as intuitive as possible. User interactions are divided into three subsequent phases. The first step simply requests input genes and a species. The second step requires the input of a secondary species, and the final step asks the user what metric to use in characterizing the output subnetworks. Visualizations of the network and the distribution of co-expression values are reported after running the first two steps. In addition, gene set enrichment is applied, and genes with over-represented GO terms can be visualized directly in the subnetwork. In the final step, we characterize the connectivity of the gene set as well as any subnetworks related to GO terms within the gene set. We typically report this as an AUROC, which specifies the degree to which the network topology allows reconstruction of the set of genes used as training, if some fraction of them are hidden (i.e., cross-validation).

Overall, an input of about 200 genes will render a network within 30 seconds and implement EGAD for GO groups within 10 seconds. An input of about 1000 genes will render a network within 5 minutes and implement EGAD for GO groups within a minute. Implementation of EGAD on the gene set takes between 30 seconds to 5 minutes depending on the selected species and the gene set to compare. In the interest of user experience, we impose an upper limit of 1000 genes since larger queries may interfere with processes of other users. For larger scale inquiries, we recommend downloading the relevant data and using *CoCoCoNetLite*, available at ftp://milton.cshl.edu/data/scripts/cococonetLite.R. We refer readers to the attached supplement(figures S5-S10) for a detailed tutorial and usage guide of CoCoCoNet.

### Downloadable

All data and R scripts used to generate results are available at ftp://milton.cshl.edu/data. Data includes gene expression networks as HDF5 files, GO annotations, gene ID conversion tables, 1-to-1 ortholog mappings, the total degree of each gene, and example gene lists. During each step, the user is also able to download relevant data. In the first two steps, co-expression networks and functional enrichment results can be downloaded, with subnetworks in coordinate format. In the final section, the user is able to download the AUROC (or AUPRC) scores of each GO term for each species.

### CASE STUDIES

#### Highly co-expressed yeast genes

Co-expression was first exploited as a global tool for characterization of gene function by Eisen et al. in a study of yeast (2), so for our first use case we returned to this original benchmark gene set to walk through a simple validational use of the main feature of CoCoCoNet. To define an interesting gene set to explore, we first pruned the Eisen list by filtering for genes with very high co-expression with at least one other gene (see supplement for details). Then, mapping this set of genes to every other species in CoCoCoNet, we see that most orthologs remain very highly co-expressed with one another, with average co-expression link-strengths above 0.8 (i.e., in the top 20%, see figure 3). Beyond the individual network links, the overall topology exhibits strikingly well-defined modules. The first cluster contains primarily ribosomal protein and translation related genes, in good agreement with group **I** in the 1998 Eisen et. al. paper. Another cluster contains predominately proteasome related genes, analogous to group **C**, while the largest cluster contains genes with functions relating to the nucleolus and translation regulation, among others. See figure S11 for a dendrogram and heatmap of these 231 genes.

**Figure 3.**
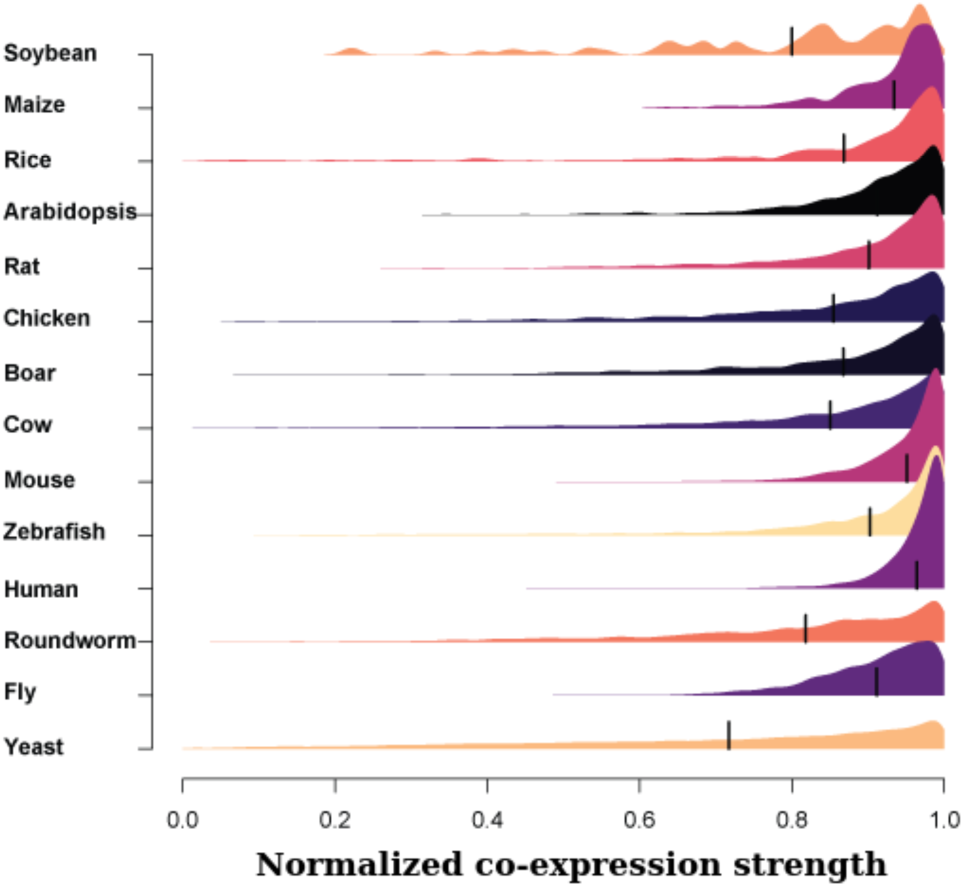
Distribution of co-expression values for ortholog mapped genes to the input of highly co-expressed yeast genes for each of the 13 other species.

Using the ortholog mapping feature of CoCoCoNet, and restricting our attention to the nucleolus (GO:0005730), we can evaluate the co-expression of the input yeast genes in other species. As expected for a structure that is common to all eukaryotes, we find that this function is highly conserved even at extreme phylogenetic distances (e.g., yeast AUROC=0.9070, Arabidopsis=0.9111, zebrafish=0.8770, fruitfly = 0.8320). A common feature of co-expression networks is hub genes which are strongly connected to many others (i.e., they have high node degree). Supporting the specificity of the nucleolus gene-gene connections, we find that our control test, which uses node degree alone to predict module connectivity, has almost no performance (AUROCs ≈0.5). Together, these results indicate that yeast nucleolus genes form a functional module that is tightly conserved across distant species (figure 4).

**Figure 4.**
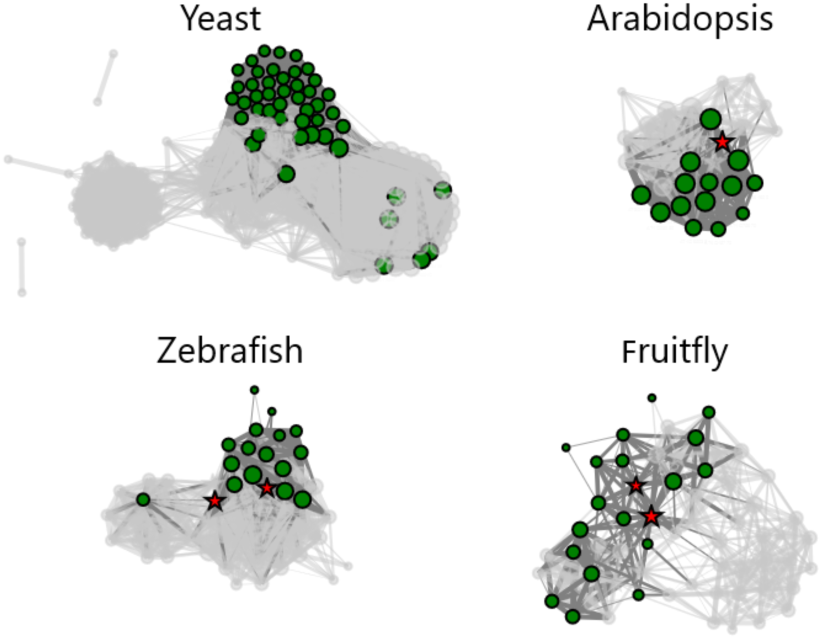
Highly co-expressed yeast (S. cerevisiae) genes are mapped to orthologous genes in Arabidopsis (A. thaliana), zebrafish (D. rerio), and fruitfly (D. melanogaster). Genes annotated with the nucleolus (GO:0005730) are highlighted, and the top 1% of connections are shown. Red stars denote highly connected genes as measured by their node degree.

#### Autism spectrum disorder associated genes

The success of translational disease research relies on the conservation of gene function between model organisms and humans. However, in many cases, it remains unclear whether disease mechanisms are sufficiently similar (40, 41). Failures of translation have been particularly notable within the neurosciences (42).

Autism spectrum disorder (ASD) is a syndrome with known phenotypic and genetic heterogeneity (43, 44, 45). Past analyses have found that ASD genes fall into two major functional categories: those involved in gene expression regulation (GER) and those involved in neuronal communication (NC) (46, 47). This suggests that cases may be subtyped based on the gene networks that are affected by rare inherited or *de novo* variants. Here, we consider the co-expression of a set of 102 genes associated with ASD identified by the Autism Sequencing Consortium in (46) along with the corresponding 1-to-1 orthologs in mouse, where functional translation is likely to be key. These genes were used as input to CoCoCoNet with default parameters.

Enrichment analyses of the 102 gene subnetworks in both mouse and human indicate that GER and NC terms are over-represented, as expected. CoCoCoNet also permits direct comparison of the GER and NC modules within- and across-species, suggesting which gene relationships can be meaningfully assessed in the mouse as a model system. Inputting the GER and NC gene sets into CoCoCoNet one at a time, we can consider the modularity of each gene set independently using the “Compute the gene set score” feature. We find that the 58 GER genes have high co-expression edge strengths with one another (average of 0.81), but they are not preferentially connected with one another at all (AUROCs of 0.415 in human and 0.556 in mouse). This suggests that while gene regulation is obviously an important function and the strong co-expression edges of the genes reflect this, they also possess equally strong relationships with other genes, making targeted translation between species difficult. In contrast, the 24 NC genes have relatively weak edge strengths (average of 0.53), but are very preferentially connected with one another (AUROCs of 0.880 in human and 0.859 in mouse), suggesting a shared mechanism that is conserved between human and mouse (figure S12).

## DISCUSSION AND OUTLOOK

Co-expression networks are useful tools for investigating gene function, but they require large-scale data aggregation to be powered, and this has limited their broader use. We have carefully curated and generated aggregate co-expression networks for 14 species, chosen because they have sufficient RNA-sequencing data as well as GO annotations. We share them via the CoCoCoNet web server to aid researchers in their comparative analyses.

CoCoCoNet provides fast enrichment and conservation scores, displayed in a user-friendly manner. Here, we have walked through two applications of CoCoCoNet, but there are many other possibilities. We make it easy to reproduce the analyses done in the web server by providing code alongside visual outputs and quantitative results. In addition, we strongly encourage users to download networks and explore them with their own biological questions in mind. We expect that future releases will encompass data from a wider variety of organisms as new research emerges.

## ACKNOWLEDGEMENTS

We are grateful to Stephan Fischer, Risa Kawaguchi, Hamsini Suresh, Sukalp Muzumdar, Shaina Lu, Benjamin Harris, Jonathan Werner, Nathan Fox, and John Hover for testing the server and offering valuable comments. We thank all researchers who make their data available.

