## Supplementary Material for "CoCoCoNet: Conserved and Comparative Co-expression Across a Diverse Set of Species"

#### Overview

Gene co-expression network analysis is a field merging network theory and transcriptomics, the uses of which are far ranging. From gene function prediction to disease gene assessment to comparative genomics, biologists use co-expression to probe a multitude of questions. However, meta-analytic approaches to build and assess transcriptional data are not common. Here, we describe our webserver for the robust meta-analysis of conserved co-expression across species. What makes CoCoCoNet unique is the number of experiments, the number of species, and the aggregation of networks. The majority of the necessary data is pre-computed so the webserver itself only requires a few seconds of compute time. The data is curated by aggregating the expression reads of many RNA-seq experiments obtained through the NCBI's SRA database (1) using the guidelines outlined in (2). The final outputs are network visualizations, comparative ranked correlation scores, and AUROCs.

#### Network aggregation pipeline

The underlying pipeline to the webserver is an *in-silico* gene-gene network building approach that involves three basic steps: co-expression network construction, network aggregation and then network assessment (**Figure S2**). The construction of co-expression networks from individual expression experiments across many samples is an established approach (3). Networks are constructed by taking gene expression profiles across samples and calculating pair-wise relationships between them. This value is typically a correlation or distance measure, and is used as the weighted edge between genes in the network (**Figure S2 A**). Co-expression is meant to reflect co-regulation, co-functionality and co-variation. However, noise is pervasive as technical confounds can be present. To overcome this, aggregation of individual networks across multiple experiments averages away noisy edges and reinforces robust links (**Figure S2 B**). Aggregation of networks built in this manner was first performed on microarray data (4) and extended to RNA-seq (2). The information content is measured by assessing the network through supervised machine learning methods that work by using the guilt-by-association principle. This is done via a neighbor-voting algorithm implemented in the EGAD Bioconductor package (5).

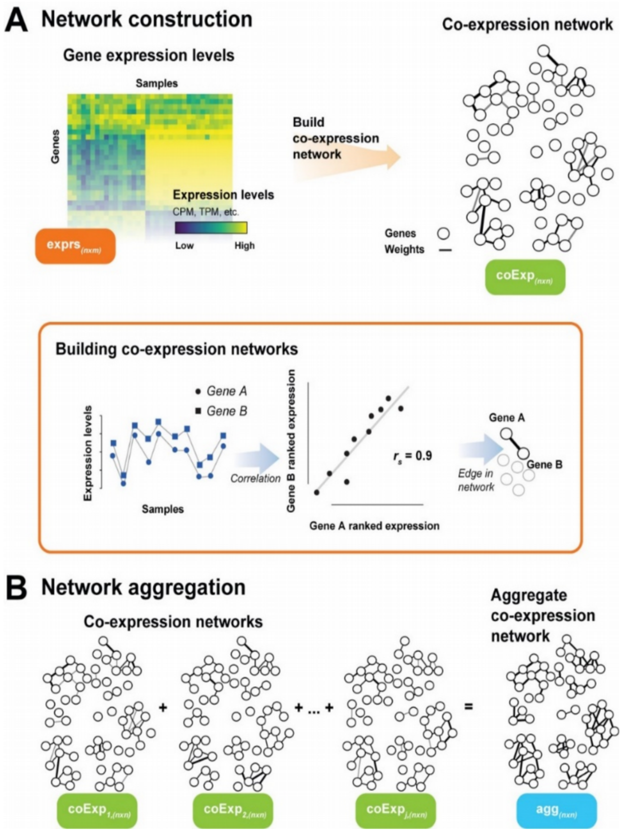

**Figure S1: Schematic of co-expression meta-analysis protocol.**
**(A)** Individual network construction involves taking a curated expression experiment with n genes and m samples and building a gene co-expression network (n by n). For each gene-gene pair, we calculate Spearman's correlation between their expression profiles, rank standardize it, and use this a weighted edged in our network. A sparsified representation is used here to distinguish the stronger correlations from the weaker ones. **(B)** For K number of networks, we aggregate by summing the edge weights and then re-rank standardizing the resulting network. Although individual networks have few strong connections, in aggregate, we robustly reconstruct modules. This is measured through the network's performance with the assessment step (not shown) which measures functional enrichment.

#### Datasets

All of the data used in the building of this webserver are listed in the supplementary tables. These include the genomes and annotation files (**Table S1**), the experiments and samples (**Table S2**) and the Gene Ontology files (**Table S3**).

**Table S1: Genome and annotation files of CoCoCoNet version 1.0**

| Species | Version of genome and annotation | File | URL |
|---|---|---|---|
| Human | GENCODE v28 | GTF | ftp://ftp.ebi.ac.uk/pub/databases/gencode/Gencode_human/release_28/ gencode.v28.chr_patch_hapl_scaff.annotation.gtf.gz |
| | GRCh38.p12 | FASTA | ftp://ftp.ebi.ac.uk/pub/databases/gencode/Gencode_human/release_31/GRCh38.p12.genome.fa.gz |
| | | Genes | ftp://ftp.ncbi.nih.gov/gene/DATA/GENE_INFO/Mammalia/Homo_sapiens.gene_info.gz<br>ftp://ftp.ncbi.nih.gov/gene/DATA/gene2ensembl.gz |
| Mouse | | GTF | ftp://ftp.ensembl.org/pub/release-94/gtf/mus_musculus/Mus_musculus.GRCm38.94.gtf.gz |
| | GRCm38 | FASTA | ftp://ftp.ensembl.org/pub/release-94/fasta/mus_musculus/dna/Mus_musculus.GRCm38.dna.primary_assembly.fa.gz |
| | | Genes | ftp://ftp.ncbi.nih.gov/gene/DATA/GENE_INFO/Mammalia/Mus_musculus.gene_info.gz<br>ftp://ftp.ncbi.nih.gov/gene/DATA/gene2ensembl.gz |
| Arabidopsis | | GTF | ftp://ftp.ensemblgenomes.org/pub/plants/release-39/gtf/arabidopsis_thaliana/Arabidopsis_thaliana.TAIR10.39.gtf.gz |
| | TAIR10 | FASTA | ftp://ftp.ensemblgenomes.org/pub/plants/release-39/fasta/arabidopsis_thaliana/dna/ Arabidopsis_thaliana.TAIR10.dna.toplevel.fa.gz |
| | | Genes | ftp://ftp.ncbi.nih.gov/gene/DATA/GENE_INFO/Plants/Arabidopsis_thaliana.gene_info.gz |
| Boar | | GTF | ftp://ftp.ensembl.org/pub/release-93/gtf/sus_scrofa/Sus_scrofa.Sscrofa11.1.93.gtf.gz |
| | Sscrofa11.1 | FASTA | ftp://ftp.ensembl.org/pub/release-93/fasta/sus_scrofa/dna/Sus_scrofa.Sscrofa11.1.dna.toplevel.fa.gz |
| | | Genes | ftp://ftp.ncbi.nih.gov/gene/DATA/GENE_INFO/Mammalia/Sus_scrofa.gene_info.gz<br>ftp://ftp.ncbi.nih.gov/gene/DATA/gene2ensembl.gz |
| Chicken | | GTF | ftp://ftp.ensembl.org/pub/release-93/gtf/gallus_gallus/Gallus_gallus.Gallus_gallus-5.0.93.gtf.gz |
| | Gallus_gallus-5.0 | FASTA | ftp://ftp.ensembl.org/pub/release-93/fasta/gallus_gallus/dna/Gallus_gallus.Gallus_gallus-5.0.dna.toplevel.fa.gz |
| | | Genes | ftp://ftp.ncbi.nih.gov/gene/DATA/GENE_INFO/Non-mammalian_vertebrates/Gallus_gallus.gene_info.gz<br>ftp://ftp.ncbi.nih.gov/gene/DATA/gene2ensembl.gz |
| Cow | | GTF | ftp://ftp.ensembl.org/pub/release-93/gtf/bos_taurus/Bos_taurus.UMD3.1.93.gtf.gz |
| | UMD3.1 | FASTA | ftp://ftp.ensembl.org/pub/release-93/fasta/bos_taurus/dna/Bos_taurus.UMD3.1.dna.toplevel.fa.gz |
| | | Genes | ftp://ftp.ncbi.nih.gov/gene/DATA/GENE_INFO/Mammalia/Bos_taurus.gene_info.gz<br>ftp://ftp.ncbi.nih.gov/gene/DATA/gene2ensembl.gz |

| | | | |
|---|---|---|---|
| Fruitfly | | GTF | ftp://ftp.ensembl.org/pub/release-93/gtf/drosophila_melanogaster/Drosophila_melanogaster.BDGP6.93.gtf.gz |
| | BDGP | FASTA | ftp://ftp.ensembl.org/pub/release-93/fasta/drosophila_melanogaster/dna/ Drosophila_melanogaster.BDGP6.dna.toplevel.fa.gz |
| | | Genes | ftp://ftp.ncbi.nih.gov/gene/DATA/GENE_INFO/Invertebrates/Drosophila_melanogaster.gene_info.gz ftp://ftp.ncbi.nih.gov/gene/DATA/gene2ensembl.gz |
| Maize | | GTF | ftp://ftp.ensemblgenomes.org/pub/plants/release-39/gtf/zea_mays/Zea_mays.AGPv4.39.gtf.gz |
| | maiAGPv4 | FASTA | ftp://ftp.ensemblgenomes.org/pub/plants/release-39/fasta/zea_mays/dna/Zea_mays.AGPv4.dna.toplevel.fa.gz |
| | | Genes | ftp://ftp.ncbi.nih.gov/gene/DATA/GENE_INFO/Plants/Zea_mays.gene_info.gz |
| Rat | | GTF | ftp://ftp.ensembl.org/pub/release-93/gtf/rattus_norvegicus/Rattus_norvegicus.Rnor_6.0.93.gtf.gz |
| | Rnor_6.0 | FASTA | ftp://ftp.ensembl.org/pub/release-93/fasta/rattus_norvegicus/dna/Rattus_norvegicus.Rnor_6.0.dna.toplevel.fa.gz |
| | | Genes | ftp://ftp.ncbi.nih.gov/gene/DATA/GENE_INFO/Mammalia/Rattus_norvegicus.gene_info.gz ftp://ftp.ncbi.nih.gov/gene/DATA/gene2ensembl.gz |
| Rice | | GTF | ftp://ftp.ensemblgenomes.org/pub/plants/release-41/gtf/oryza_sativa/Oryza_sativa.IRGSP-1.0.41.gtf.gz |
| | rIRGSP-1.0 | FASTA | ftp://ftp.ensemblgenomes.org/pub/plants/release-41/fasta/oryza_sativa/dna/Oryza_sativa.IRGSP-1.0.dna.toplevel.fa.gz |
| | | Genes | ftp://ftp.ncbi.nih.gov/gene/DATA/GENE_INFO/Plants/Oryza_sativa.gene_info.gz |
| Roundworm | | GTF | ftp://ftp.ensembl.org/pub/release-94/gtf/caenorhabditis_elegans/Caenorhabditis_elegans.WBcel235.94.gtf.gz |
| | WBcel235 | FASTA | ftp://ftp.ensembl.org/pub/release-94/fasta/caenorhabditis_elegans/dna/ Caenorhabditis_elegans.WBcel235.dna.toplevel.fa.gz |
| | | Genes | ftp://ftp.ncbi.nih.gov/gene/DATA/GENE_INFO/Invertebrates/Caenorhabditis_elegans.gene_info.gz ftp://ftp.ncbi.nih.gov/gene/DATA/gene2ensembl.gz |
| Soybean | | GTF | ftp://ftp.ensemblgenomes.org/pub/plants/release-41/gtf/glycine_max/Glycine_max.Glycine_max_v2.0.41.gtf.gz |
| | Glycine_max_v2.0 | FASTA | ftp://ftp.ensemblgenomes.org/pub/plants/release-41/fasta/glycine_max/dna/ Glycine_max.Glycine_max_v2.0.dna.toplevel.fa.gz |
| | | Genes | ftp://ftp.ncbi.nih.gov/gene/DATA/gene_info.gz |
| Yeast | | GTF | ftp://ftp.ensembl.org/pub/release-93/gtf/saccharomyces_cerevisiae/Saccharomyces_cerevisiae.R64-1-1.93.gtf.gz ftp://ftp.ncbi.nih.gov/gene/DATA/gene2ensembl.gz |
| | R64-1-1 | FASTA | ftp://ftp.ensembl.org/pub/release-93/fasta/saccharomyces_cerevisiae/dna/Saccharomyces_cerevisiae.R64-1-1.dna.toplevel.fa.gz |
| | | Genes | ftp://ftp.ncbi.nih.gov/gene/DATA/GENE_INFO/Fungi/Saccharomyces_cerevisiae.gene_info.gz |
| Zebrafish | | GTF | ftp://ftp.ensembl.org/pub/release-93/gtf/danio_rerio/Danio_rerio.GRCz11.93.chr_patch_hapl_scaff.gtf.gz |
| | GRCz11 | FASTA | ftp://ftp.ensembl.org/pub/release-93/fasta/danio_rerio/dna/Danio_rerio.GRCz11.dna.primary_assembly.fa.gz |
| | | Genes | ftp://ftp.ncbi.nih.gov/gene/DATA/GENE_INFO/Non-mammalian_vertebrates/Danio_rerio.gene_info.gz ftp://ftp.ncbi.nih.gov/gene/DATA/gene2ensembl.gz |

**Table S2: List of experiments used per species**

| Species | Scientific name | NCBI taxon ID | Link |
|---|---|---|---|
| **Human** | *Homo sapiens* | 9606 | ftp://milton.cshl.edu/data/summaryFiles/human_summary.csv |
| **Mouse** | *Mus musculus* | 10090 | ftp://milton.cshl.edu/data/summaryFiles/mouse_summary.csv |
| **Arabidopsis** | *Arabidopsis thaliana* | 3702 | ftp://milton.cshl.edu/data/summaryFiles/arabidopsis_summary.csv |
| **Boar** | *Sus scrofa* | 9823 | ftp://milton.cshl.edu/data/summaryFiles/boar_summary.csv |
| **Chicken** | *Gallus gallus* | 9031 | ftp://milton.cshl.edu/data/summaryFiles/chicken_summary.csv |
| **Cow** | *Bos taurus* | 9913 | ftp://milton.cshl.edu/data/summaryFiles/cow_summary.csv |
| **Fruitfly** | *Drosophila melanogaster* | 7227 | ftp://milton.cshl.edu/data/summaryFiles/fruitfly_summary.csv |
| **Maize** | *Zea mays* | 4577 | ftp://milton.cshl.edu/data/summaryFiles/maize_summary.csv |
| **Rat** | *Rattus norvegicus* | 10116 | ftp://milton.cshl.edu/data/summaryFiles/rat_summary.csv |
| **Rice** | *Oryza sativa* | 39947 | ftp://milton.cshl.edu/data/summaryFiles/rice_summary.csv |
| **Roundworm** | *Caenorhabditis elegans* | 6239 | ftp://milton.cshl.edu/data/summaryFiles/roundworm_summary.csv |
| **Soybean** | *Glycine max* | 3847 | ftp://milton.cshl.edu/data/summaryFiles/soybean_summary.csv |
| **Yeast** | *Saccharomyces cerevisiae* | 559292 | ftp://milton.cshl.edu/data/summaryFiles/yeast_summary.csv |
| **Zebrafish** | *Danio rerio* | 7955 | ftp://milton.cshl.edu/data/summaryFiles/zebrafish_summary.csv |

**Table S3: Gene ontology files for CoCoCoNet version 1.0 obtained by merging data from NCBI and biomaRt with the exception of maize**

| Species | Scientific name | NCBI taxon ID | Gene annotation file |
|---|---|---|---|
| Human | *Homo sapiens* | 9606 | https://ftp.ncbi.nlm.nih.gov/gene/DATA/gene2go.gz |
| Mouse | *Mus musculus* | 10090 | https://ftp.ncbi.nlm.nih.gov/gene/DATA/gene2go.gz |
| Arabidopsis | *Arabidopsis thaliana* | 3702 | https://ftp.ncbi.nlm.nih.gov/gene/DATA/gene2go.gz |
| Boar | *Sus scrofa* | 9823 | https://ftp.ncbi.nlm.nih.gov/gene/DATA/gene2go.gz |
| Chicken | *Gallus gallus* | 9031 | https://ftp.ncbi.nlm.nih.gov/gene/DATA/gene2go.gz |
| Cow | *Bos taurus* | 9913 | https://ftp.ncbi.nlm.nih.gov/gene/DATA/gene2go.gz |
| Fruitfly | *Drosophila melanogaster* | 7227 | https://ftp.ncbi.nlm.nih.gov/gene/DATA/gene2go.gz |
| Maize | *Zea mays* | 4577 | biomaRt only |
| Rat | *Rattus norvegicus* | 10116 | https://ftp.ncbi.nlm.nih.gov/gene/DATA/gene2go.gz |
| Rice | *Oryza sativa* | 39947 | https://ftp.ncbi.nlm.nih.gov/gene/DATA/gene2go.gz |
| Roundworm | *Caenorhabditis elegans* | 6239 | https://ftp.ncbi.nlm.nih.gov/gene/DATA/gene2go.gz |
| Soybean | *Glycine max* | 3847 | https://ftp.ncbi.nlm.nih.gov/gene/DATA/gene2go.gz |
| Yeast | *Saccharomyces cerevisiae* | 559292 | https://ftp.ncbi.nlm.nih.gov/gene/DATA/gene2go.gz |
| Zebrafish | *Danio rerio* | 7955 | https://ftp.ncbi.nlm.nih.gov/gene/DATA/gene2go.gz |

#### Experiment metadata and network performances

The total number of experiments used per species and the total number of samples are shown in **Table S4**. The properties of these experiments per species are shown as distributions in **Figure S2** (proportion of missing genes), **Figure S3** (correlations to local expression) and **Figure S4** (correlations to global expression). In addition, we ran EGAD on the aggregate networks with their own GO annotations, using either the high confidence set of genes, or almost all genes. The performance AUROCs are listed in **Table S4**, along with the number of genes and GO terms. Scores range between 0.64 and 0.76, with mean 0.71 for the high confidence networks, and 0.68 for all genes. Finally, we show the number of orthologs between species in **Table S5**.

**Table S4: Overview of data counts and network performances.**

| Species | Experiments | Samples | Illumina HiSeq 2000/2500 samples | All genes | High confidence genes | GO terms | Aggregate (high confidence) network performance (AUROC) | Aggregate (all genes) network performance (AUROC) | Protein coding / ncRNA (high confidence) | Protein coding / ncRNA (all genes) |
|---|---|---|---|---|---|---|---|---|---|---|
| **Arabidopsis** | 124 | 3,154 | 2,397 | 34,262 | 20,879 | 10,615 | 0.7086 | 0.6792 | 20,416 / 12 | 27,407 / 132 |
| **Boar** | 49 | 1,772 | 1,563 | 25,880 | 14,142 | 20,470 | 0.7000 | 0.6805 | 13,321 / 58 | 17,840 / 390 |
| **Chicken** | 42 | 1,272 | 1,076 | 24,881 | 14,059 | 19,631 | 0.6859 | 0.6526 | 11,872 / 43 | 14,431 / 729 |
| **Cow** | 49 | 2,187 | 1,598 | 24,616 | 14,237 | 20,671 | 0.6993 | 0.6829 | 13,212 / 22 | 17,253 / 604 |
| **Fruitfly** | 89 | 2,734 | 2,297 | 17,737 | 11,529 | 12,540 | 0.7333 | 0.7158 | 10,898 / 544 | 13,889 / 3,036 |
| **Human** | 90 | 7,075 | 6,698 | 64,485 | 24,631 | 22,517 | 0.6769 | 0.6395 | 15,982 / 1,650 | 19,266 / 5,743 |
| **Maize** | 76 | 2,952 | 2,562 | 46,272 | 25,981 | 8,470 | 0.7176 | 0.6763 | 16,535 / 1 | 19,284 / 2 |
| **Mouse** | 85 | 3,359 | 2,759 | 54,446 | 21,377 | 22,833 | 0.7187 | 0.6991 | 16,571 / 1,591 | 22,662 / 6,550 |
| **Rat** | 68 | 1,892 | 1,500 | 32,883 | 15,178 | 22,718 | 0.6969 | 0.6777 | 14,099 / 37 | 20,190 / 625 |
| **Rice** | 23 | 1,192 | 1,119 | 36,850 | 16,062 | 8,705 | 0.7080 | 0.6679 | 11,528 / 0 | 18,641 / 0 |
| **Roundworm** | 27 | 782 | 670 | 46,778 | 11,995 | 10,188 | 0.7339 | 0.6484 | 11,686 / 57 | 19,800 / 24,031 |
| **Soybean** | 25 | 578 | 517 | 58,994 | 29,165 | 7,829 | 0.7231 | 0.6638 | 25,598 / 10 | 39,234 / 43 |
| **Yeast** | 65 | 2,690 | 2,354 | 7,126 | 6,072 | 9,289 | 0.7355 | 0.7578 | 5,640 / 82 | 5,819 / 96 |
| **Zebrafish** | 83 | 7,878 | 7,619 | 32,520 | 20,890 | 13,809 | 0.7158 | 0.7041 | 19,351 / 187 | 22,662 / 709 |
| **Total** | **895** | **39,517** | **34,729** | | | | | | | |

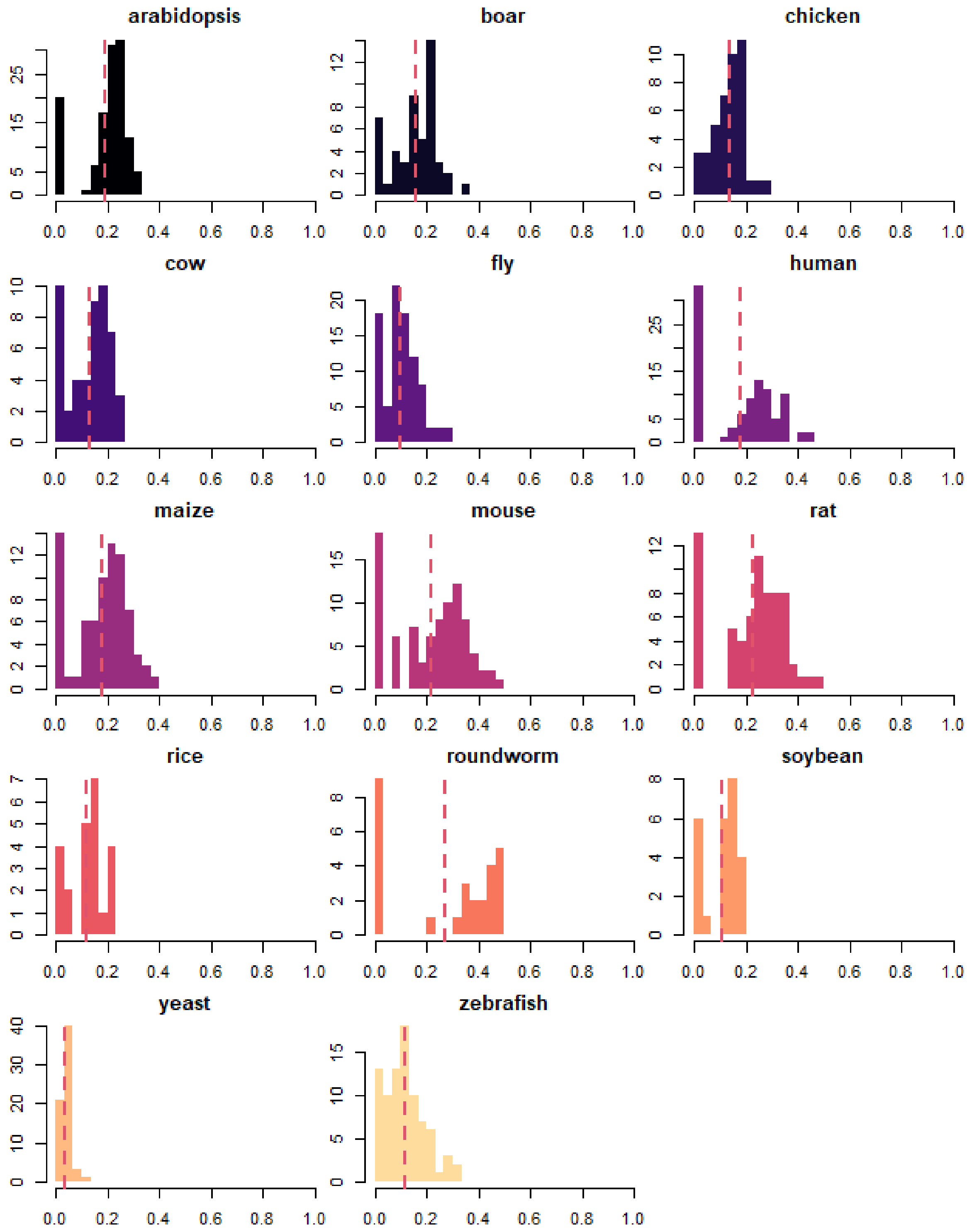

**Figure S2: Distribution of the proportion of genes missing per experiment for each species.**
Horizontal axes shows the proportion of genes missing, while vertical axes depict the number of experiments for each species. Dashed red lines correspond to the averages.

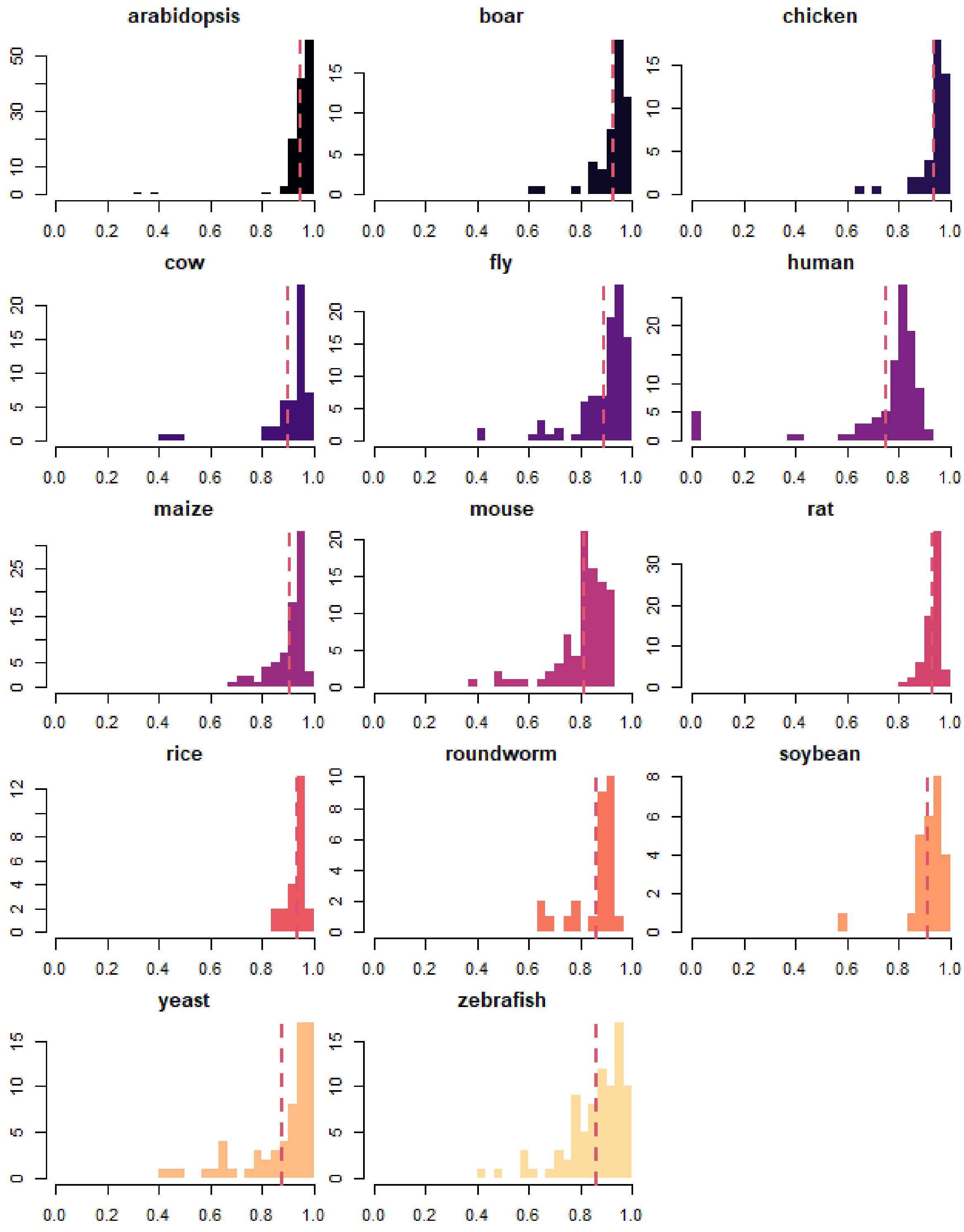

**Figure S3: Distribution of correlations to average expression within an experiment per species.** Horizontal axes shows the Spearman's correlation to the *average expression within an experiment*, while the vertical axes depict the number of experiments for each species. Dashed red lines correspond to the averages.

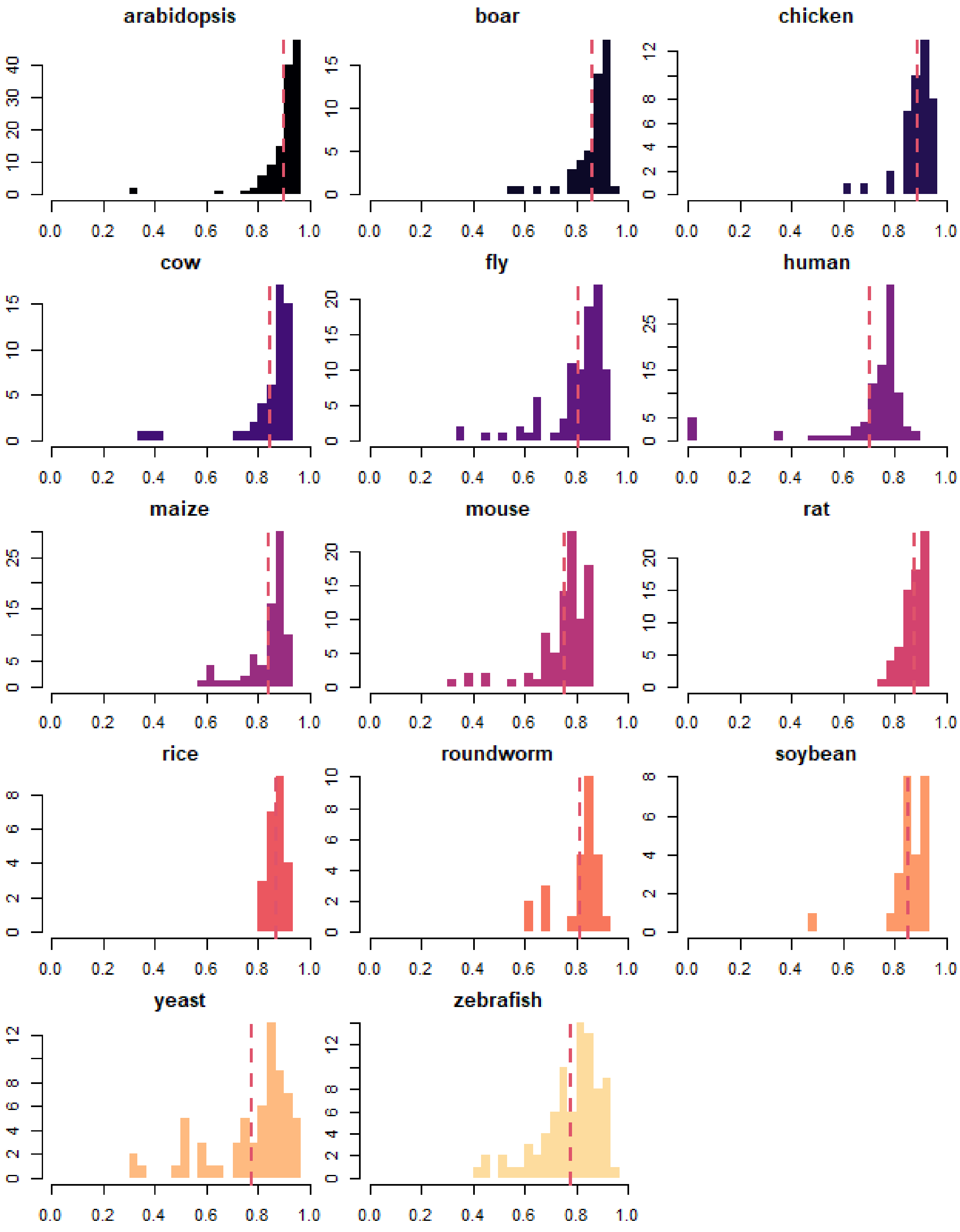

**Figure S4: Distribution of correlations to the global average expression for each experiment per species.** Horizontal axes shows the Spearman's correlation to the *global average expression*, while the vertical axis depicts the number of experiments. The global average expression is the average of *within experiment average expression*. Dashed red lines correspond to the averages.

**Table S5: Number of 1-to-1 orthologs (genes) between species**

| | Arabidopsis | Boar | Chicken | Cow | Fruitfly | Human | Maize | Mouse | Rat | Rice | Roundworm | Soybean | Yeast | Zebrafish |
|---|---|---|---|---|---|---|---|---|---|---|---|---|---|---|
| **Arabidopsis** | - | 1054 | 1002 | 1058 | 988 | 987 | 3346 | 1038 | 1008 | 4076 | 834 | **1626** | 803 | 985 |
| **Boar** | 1054 | - | 7473 | **14900** | 2342 | 11976 | 858 | 12452 | 12343 | 1103 | 1837 | 416 | 1211 | 5118 |
| **Chicken** | 1002 | 7473 | - | 7493 | 2237 | 6941 | 808 | 7277 | 7168 | 1047 | 1758 | 382 | 1158 | 4784 |
| **Cow** | 1058 | **14900** | **7493** | - | **2344** | 11953 | 850 | 12410 | 12311 | 1116 | 1838 | 404 | 1205 | **5136** |
| **Fruitfly** | 988 | 2342 | 2237 | 2344 | - | 2167 | 776 | 2310 | 2181 | 1035 | **2190** | 350 | **1265** | 2074 |
| **Human** | 987 | 11976 | 6941 | 11953 | 2167 | - | 802 | 12742 | 12538 | 1053 | 1724 | 378 | 1144 | 4780 |
| **Maize** | 3346 | 858 | 808 | 850 | 776 | 802 | - | 842 | 807 | **7654** | 651 | 1331 | 657 | 790 |
| **Mouse** | 1038 | 12452 | 7277 | 12410 | 2310 | **12742** | 842 | - | **14739** | 1093 | 1798 | 397 | 1193 | 5007 |
| **Rat** | 1008 | 12343 | 7168 | 12311 | 2181 | 12538 | 807 | **14739** | - | 1053 | 1702 | 387 | 1145 | 4913 |
| **Rice** | **4076** | 1103 | 1047 | 1116 | 1035 | 1053 | **7654** | 1093 | 1053 | - | 888 | 1468 | 864 | 1019 |
| **Roundworm** | 834 | 1837 | 1758 | 1838 | 2190 | 1724 | 651 | 1798 | 1702 | 888 | - | 280 | 1188 | 1615 |
| **Soybean** | 1626 | 416 | 382 | 404 | 350 | 378 | 1331 | 397 | 387 | 1468 | 280 | - | 259 | 357 |
| **Yeast** | 803 | 1211 | 1158 | 1205 | 1265 | 1144 | 657 | 1193 | 1145 | 864 | 1188 | 259 | - | 1109 |
| **Zebrafish** | 985 | 5118 | 4784 | 5136 | 2074 | 4780 | 790 | 5007 | 4913 | 1019 | 1615 | 357 | 1109 | - |

Bold terms indicate the highest overlap (by column).

### Tutorial

To use CoCoCoNet (**Figure S5**), simply input a list of genes (or a single gene) as gene symbols and the corresponding species to be used in the construction of the network. Given these, you can select optional parameters (described below), generate the results and view the distribution of co-expression values as well as the network. CoCoCoNet also allows you to compare the network with another species using a 1-to-1 ortholog mapping (6) as well as perform a GBA analysis using EGAD (5).

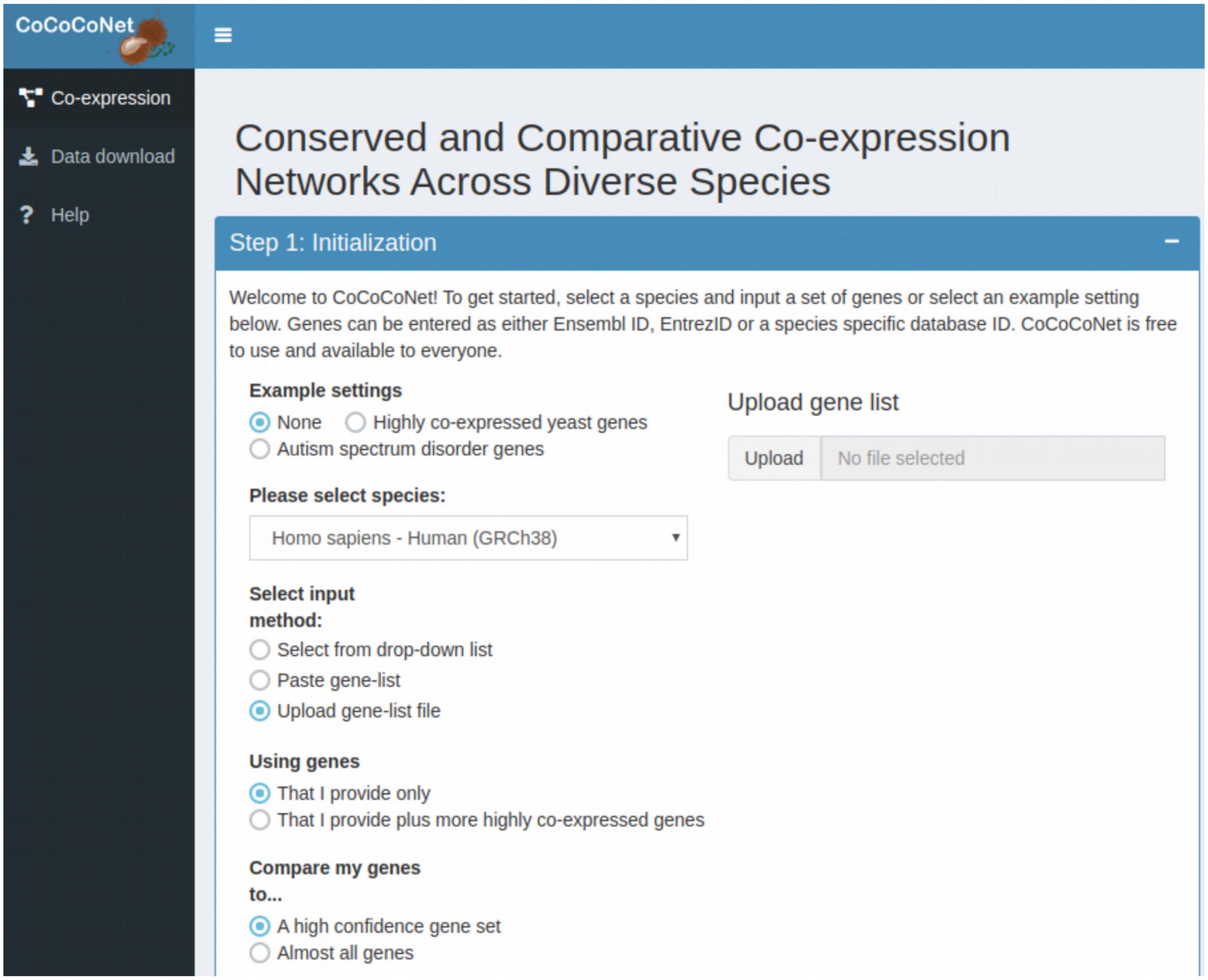

**Figure S5: Screenshot of homepage of CoCoCoNet version 1.0.**

#### Initialization

Preloaded genes can be used to test the server's functionality with the **"Example settings"**. Users can choose either the top 231 co-expressed yeast genes studied by Eisen et al (7) or the top 102 co-expressed genes associated with Autism Spectrum Disorder (ASD) from Satterstrom et al (8). More on these examples can be found in the use cases section of the manuscript and this document. In the steps below, we have selected the yeast set (**Figure S6**).

**"Select Input method:"** allows users to choose how to load in a gene list. Currently, users can select genes from a drop-down list, paste a comma separated list, or upload a file with genes listed in new lines. Toggling between these will change the adjacent **"Upload gene list"** box.

Users can extend their analyses beyond their gene set with the **"Using genes"** options.

- **"That I provide only"** will construct the network using only selected genes.
- **"That I provide plus more highly co-expressed genes"** will select the most closely related genes not in the provided set. Here we define the "relation" as having the largest weighted degree of edges connected to the provided genes.

**"Compare my genes to"** options allow users to limit the genes to a high confidence set or all genes based on expression levels.

- **"A high confidence gene set"** will match input genes to only a subset of genes filtered on minimum expression level across experiments.
- **"Almost all genes"** will match your genes to a very lightly filtered gene set.

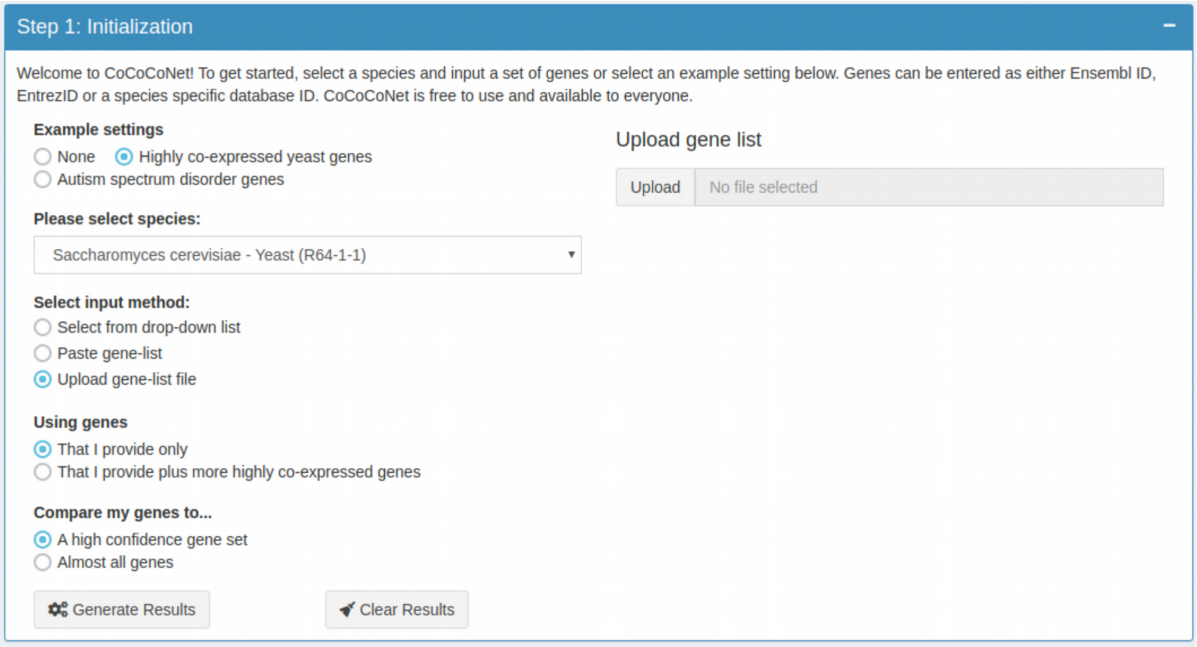

**Figure S6: Screenshot of gene input step.**
Selecting the **"Highly co-expressed yeast genes"** loads the genes onto the server, and automatically selects the species (here yeast).

#### Visualizing results: network and co-expression values of gene set

Once your genes have been properly loaded, the **"Generate Results"** and **"Clear Results"** options will appear. **Selecting "Generate Results"** will display the network, the distribution of co-expression values, and a sliding threshold bar that allows the user to filter the network to include only connections greater than this threshold. The user also has the option to highlight genes along with their direct connections or highlight genes with specified enriched GO terms. Using the example setting of **"Highly co-expressed yeast genes"** and selecting genes with GO term **"ribosome"** from the drop-down box gives the following network – highlighting genes involved in the ribosome (**Figure S7**).

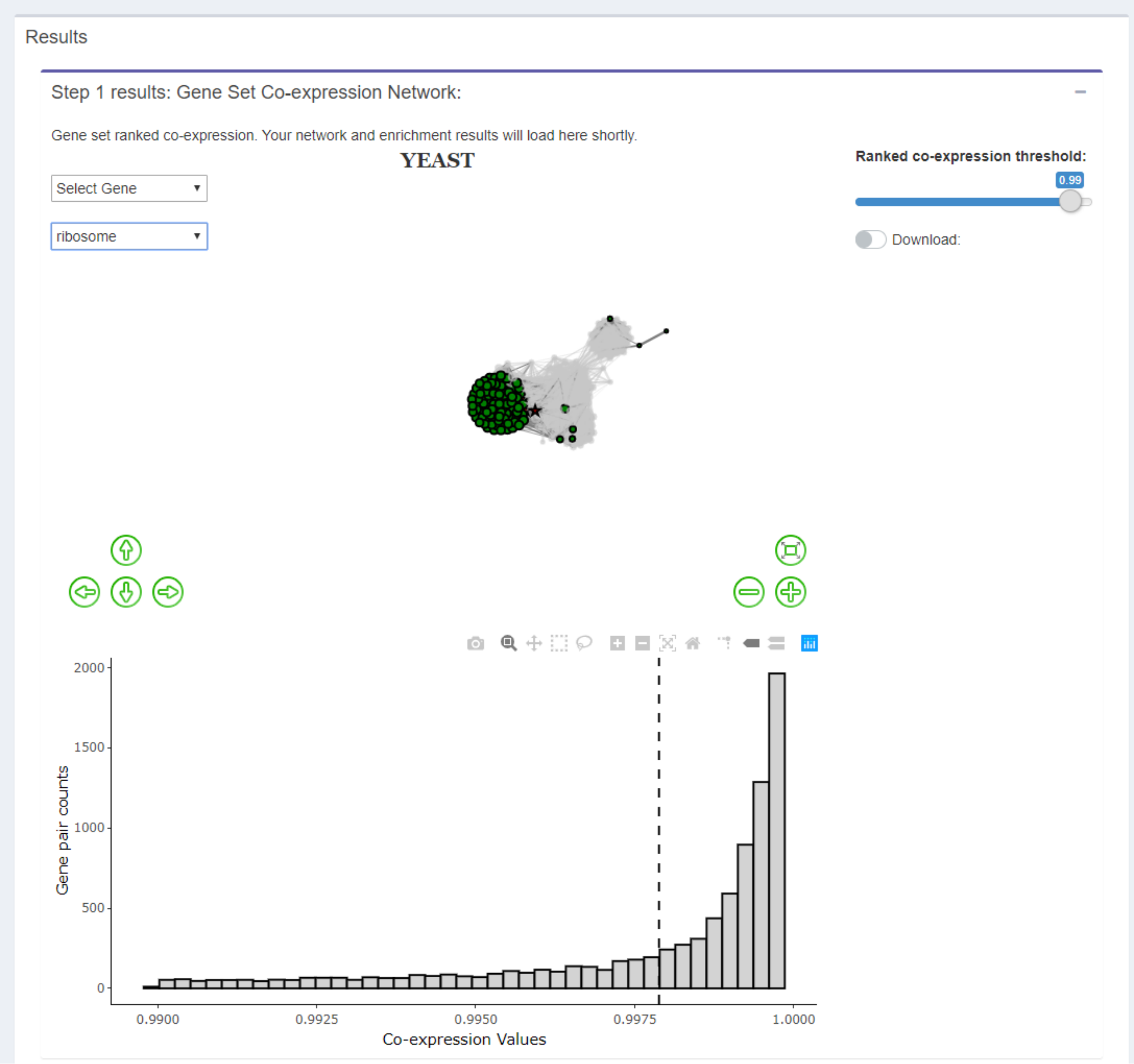

**Figure S7: Screenshot of results step 1.**
Highly co-expressed yeast genes network highlighting the ribosomal genes.

#### Exporting results

User results can also be downloaded by toggling the **"Download"** option at each stage. Here, the user can download the gene list, gene pairs and respective co-expression, above the defined threshold, GO enrichment result, and a list of genes with their enriched GO term (**Figure S8**). The network can be exported by right-clicking and saving the image and the histogram can be exported by selecting the plotly download button.

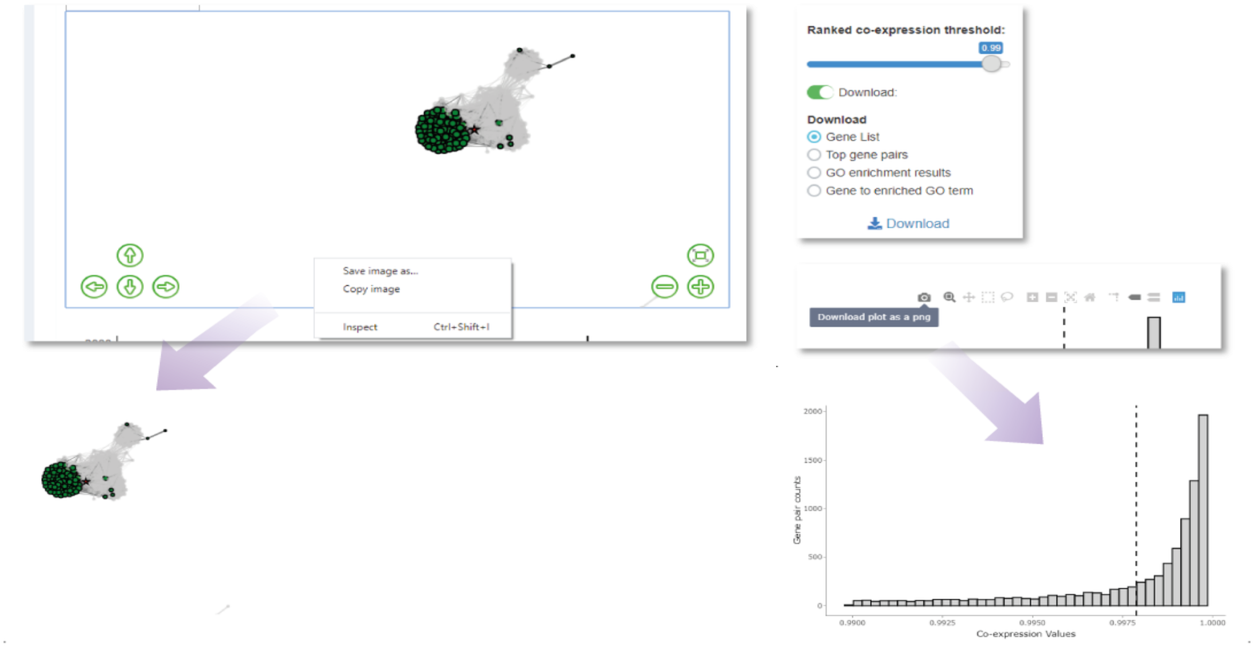

**Figure S8: Screenshots of all download options and their outputs below.**
Download box in top right corner, right-clicking on network image, or hovering over histogram for plotly options.

#### Conserved species comparison

The next section allows the user to select a second species to compare to the first. Doing so will select genes of the second species with a 1-to-1 ortholog of the provided gene set. From the drop-down menu, select a second species. After selection, "**Generate**" and "**Clear Section**" buttons will appear where selecting **"Generate"** will again display the network, the co-expression value distribution, and a sliding threshold bar. Again, the user has the option to highlight genes along with its nearest neighbors or highlight genes by GO term. Here we've selected *Psmb4, proteasome subunit, beta type 4* (ENSMUSG00000005779) as an example. Results can be exported as in the previous section (**Figure S9)**.

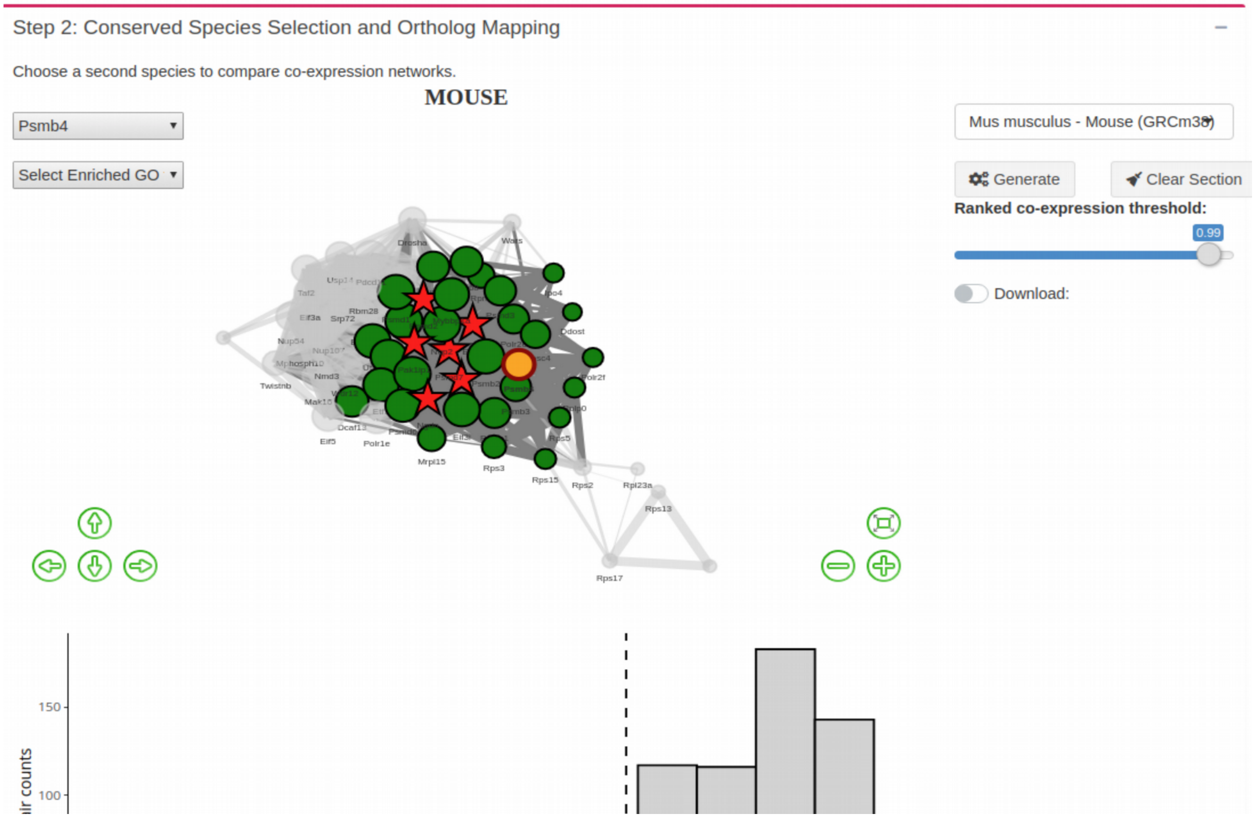

**Figure S9: Screenshot of species selection step.**

#### Comparative assessment using EGAD

The final section allows the user to perform Guilt by Association (GBA) analysis on the input genes and the corresponding 1-to-1 ortholog using EGAD (5). EGAD analyzes enriched GO terms of each species using either neighbor voting or by node degree and reports the corresponding area under that receiver operating curve (AUROC) or the precision recall curve (AURPC) across 3 cross-validation folds (**Figure S10**). Results can be exported as in the previous sections.

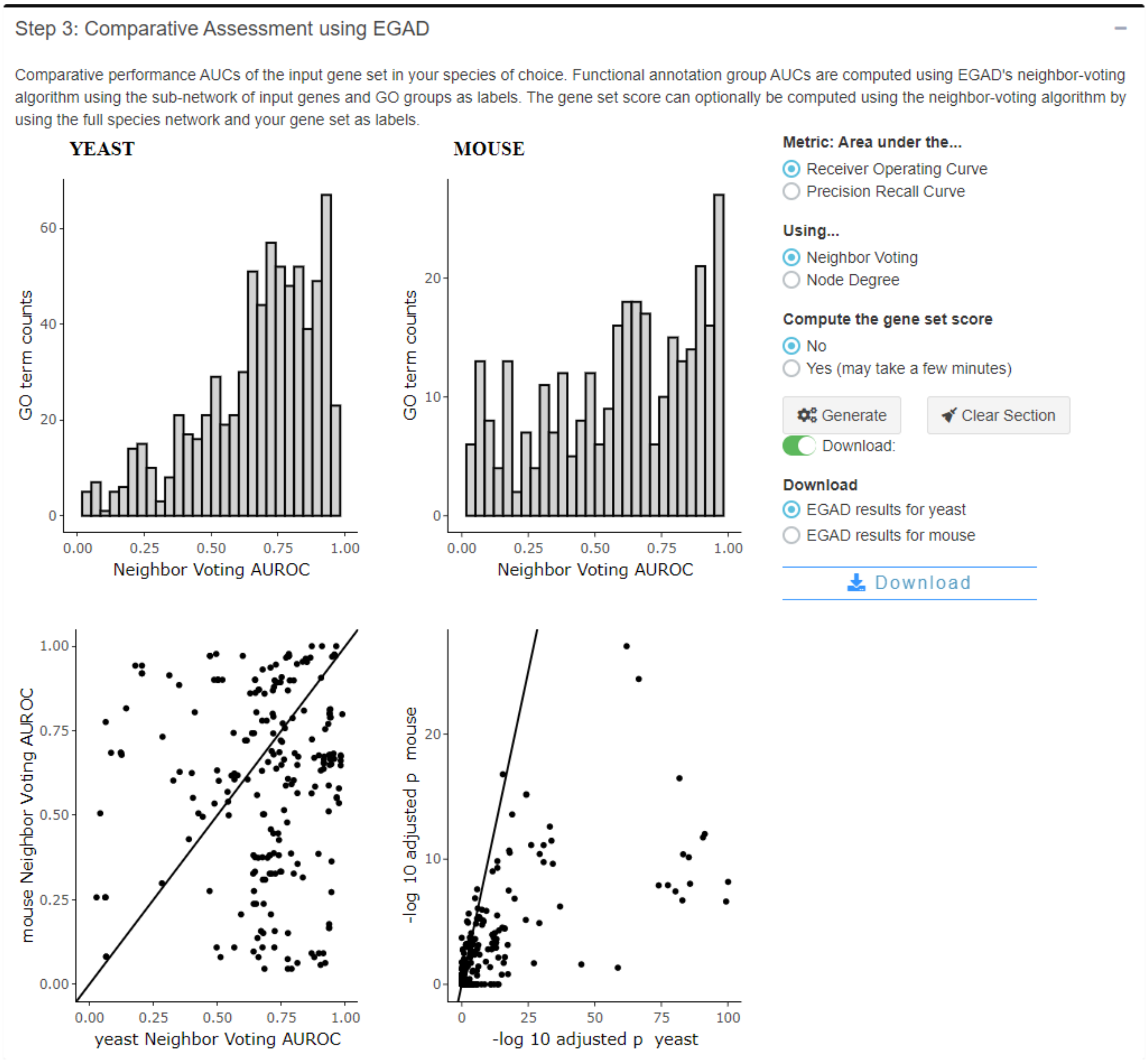

**Figure S10: Screenshot of comparative assessment step.**
Here we've run EGAD, displaying results as AUROCs and GO enrichment adjusted p-values.

### Case studies

#### Highly co-expressed yeast genes

Genes were downloaded from the supplementary files of Eisen et al (7) (https://www.pnas.org/highwire/filestream/584765/field_highwire_adjunct_files/1/3917data.xls). Of these genes, we looked to those highly co-expressed (top 0.03%) within the dataset resulting in 231 genes. Extending this example, we can take this gene set and export all co-expression values between these genes using CoCoCoNet, building a dense co-expression network (**Figure S11 A**)**.** We can then look at the values of correlations of these genes in all other species, showing on average that the 1-to-1 orthologs have highly conserved co-expression (**Figure S11 B**)**,** with mean co-expression all above 0.8. We show the subset of conserved modules in a few species in the main text.

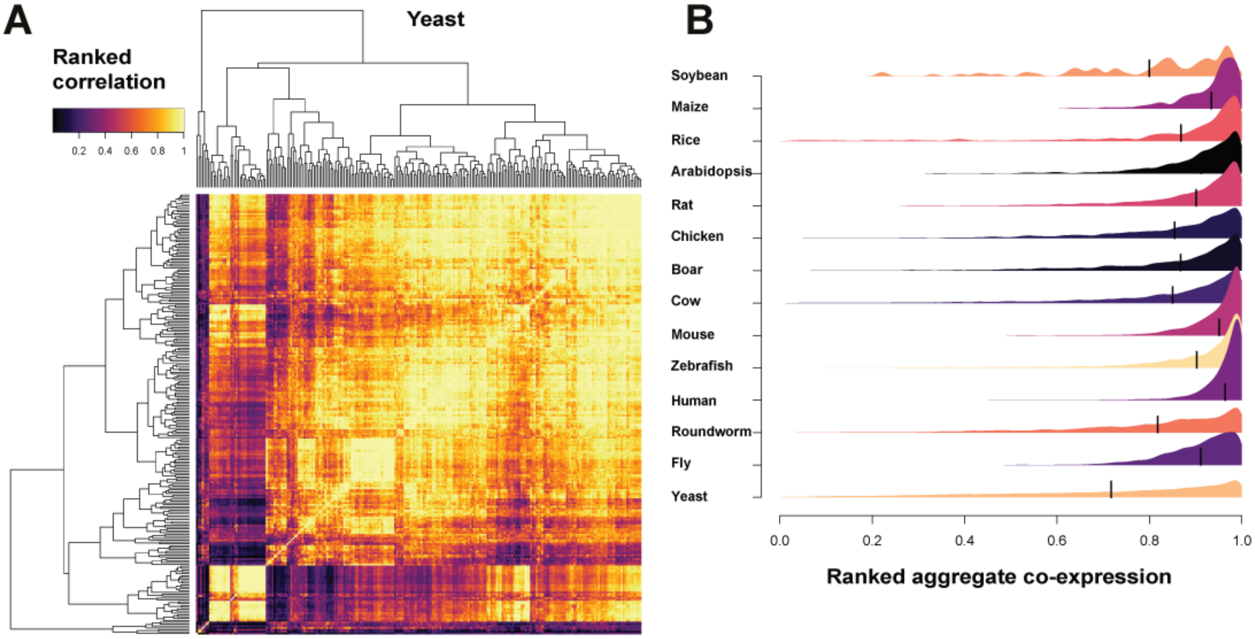

**Figure S11: Yeast co-expression of 231 yeast gene pairs.**
(A) Co-expression network as a matrix. (B) Distributions of correlations in others species ordered by number of orthologs. No threshold on the ranked correlations. Most orthologs are highly co-expressed on average.

#### Autism spectrum disorder associated genes

A recent study on autism using whole exomes was conducted by Satterstrom et al (8). We took their 102 candidates from the supplementary material (Table S2) and imported them into CoCoCoNet as the ASD example set. We compared their co-expression to those in the mouse, which is used as a model organism for many ASD studies. Of these 102 genes, Satterstrom et al (8) identified 24 as neuronal communication (NC) related while another 58 were identified as gene expression regulatory (GER). Inputting these two sets into CoCoCoNet separately and computing the "Gene set score" shows that NC genes are highly connected (AUROCs > 0.8) whereas GER genes are not (AURCOs ~ 0.5). See **figure S12**.

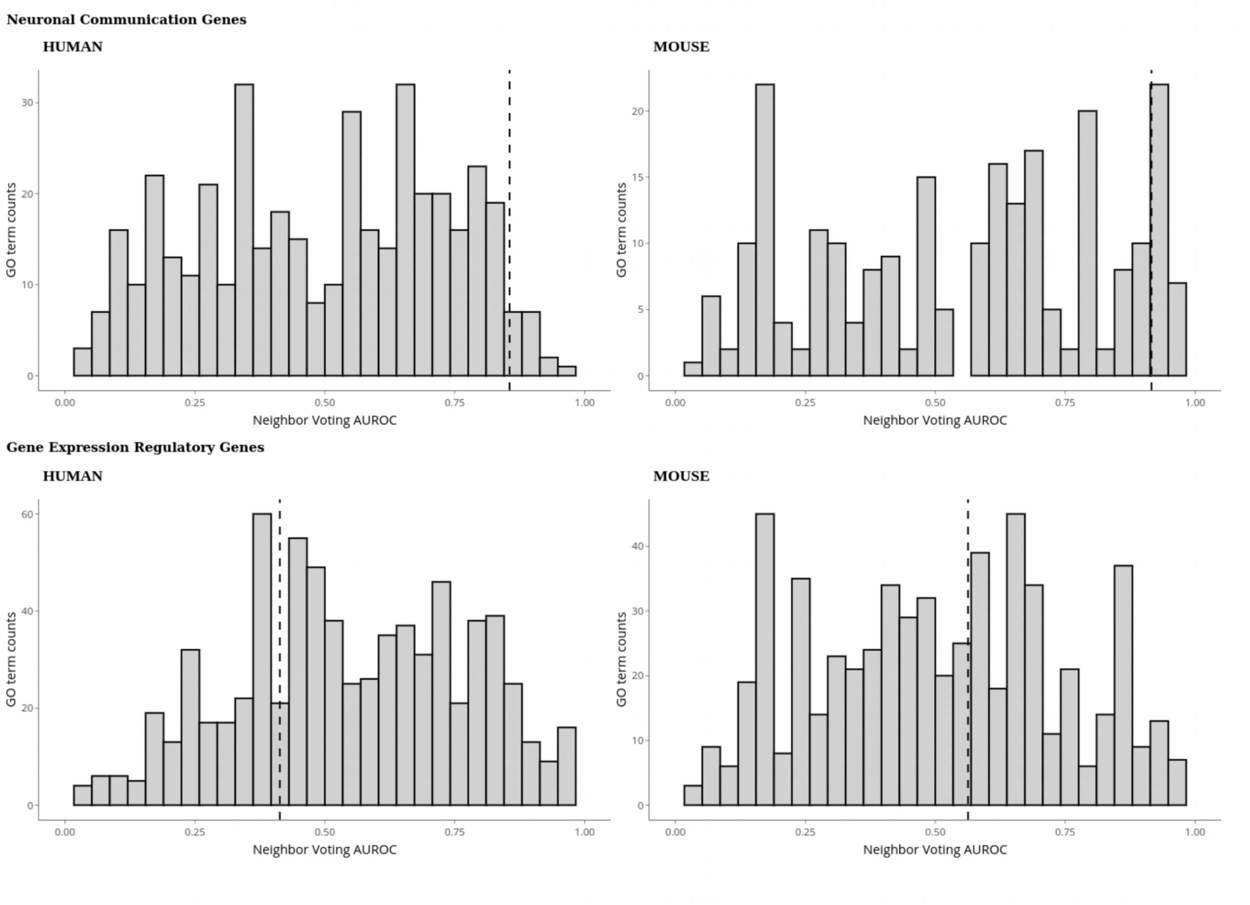

**Figure S12: Comparison of neuronal communication genes and gene expression regulatory genes.** Top panels show the GO term AUROC scores for networks limited to the 24 NC genes in human along with the corresponding 1-to-1 ortholog in mouse. These genes are very preferentially connected with AUROCs greater than 0.8 for both species (vertical dashed lines). Bottom panels show the same but for a network limited to the 58 GER genes. These genes are *not* preferentially connected with AUROCs between 0.4 and 0.6 (vertical dashed lines).
